# Whole-brain functional connectivity predicts regional tau PET in preclinical Alzheimer’s disease

**DOI:** 10.1101/2024.04.02.587791

**Authors:** Hamid Abuwarda, Anne Trainer, Corey Horien, Xilin Shen, Suyeon Ju, R. Todd Constable, Carolyn Fredericks

## Abstract

Preclinical Alzheimer’s disease (AD), characterized by the abnormal accumulation of amyloid prior to cognitive symptoms, presents a critical opportunity for early intervention. Past work has described functional connectivity changes in preclinical disease, yet the interplay between AD pathology and the functional connectome during this window remains unexplored.

We applied connectome-based predictive modeling to investigate the ability of resting-state whole-brain functional connectivity to predict tau (18F-flortaucipir) and amyloid (18F-florbetapir) PET binding in a preclinical AD cohort (A4, *n*=342, age 65-85). Separate predictive models were developed for each of 14 regions, and model performance was assessed using a Spearman’s correlation between predicted and observed PET binding standard uptake value ratios. We assessed the validity of significant models by applying them to an external dataset, and visualized the underlying connectivity that was positively and negatively correlated to posterior cingulate tau binding, the most successful model.

We found that whole brain functional connectivity predicts regional tau PET, outperforming amyloid PET models. The best performing tau models were for regions affected in Braak stage IV-V regions (posterior cingulate, precuneus, lateral occipital cortex, middle temporal, inferior temporal, and Bank STS), while models for regions of earlier tau pathology (entorhinal, parahippocampal, fusiform, and amygdala) performed poorly. Importantly, tau models generalized to a symptomatic AD cohort (ADNI; amyloid positive, *n* = 211, age 55-90), in tau-elevated but not tau-negative individuals. For the posterior cingulate A4 tau model, the most successful model, the predictive edges positively correlated with posterior cingulate tau predominantly came from nodes within temporal, limbic, and cerebellar regions. The most predictive edges negatively associated to tau were from nodes of heteromodal association areas, particularly within the prefrontal and parietal cortices.

These findings reveal that whole-brain functional connectivity predicts tau PET in preclinical AD and generalizes to a clinical dataset specifically in individuals with abnormal tau PET, highlighting the relevance of the functional connectome for the early detection and monitoring of AD pathology.

## Introduction

Alzheimer’s disease (AD) is a progressive neurodegenerative process characterized by the buildup of amyloid and tau pathology. Amyloid spreads quickly and diffusely throughout the neocortex, a process which can begin decades prior to symptom onset^10,11^. In amnestic AD, tau spread parallels clinical symptomatology and occurs in a stereotyped manner, following Braak staging^12^. Cortical tau buildup typically begins in the entorhinal cortex, then spreads to the medial and lateral temporal cortices and cingulate gyrus, corresponding with the first overt cognitive changes. The later stages of the disease are subsequently characterized by tau spread to the remaining neocortex, when AD dementia begins^12–14^.

AD is also associated with characteristic changes in functional connectivity. The default mode network (DMN), which is associated with short-term memory function and self-referential processing, is specifically affected in amnestic AD^15–18^. Preclinical disease is also characterized by regional connectivity changes in the DMN and early Braak stage regions^19,20^. However, while correlation- and regression-based studies that focus exclusively on these regions and networks are insightful and can advance our knowledge of the disease, they may also overlook important features of the disease due to the limited scope of analysis.

Importantly, the relationship between preclinical AD pathology and associated changes in the functional connectome is not well known. Evidence from animal models and emerging evidence from human studies suggest amyloid-beta induces a state of hyperexcitability^1–4^and hyperconnectivity^5,6^. Amyloid deposition has been noted to overlap significantly with the DMN especially in the very early stages of AD. On the other hand, tau appears to travel along paths of functional connectivity^7–9^. To further our understanding of this relationship, an unbiased, data-driven approach is needed. Predictive models that consider features of the entire functional connectome can help fill these gaps, especially due to their ability to minimize in sample inflation through a cross-validation procedure^21,22^, crucial for studies with potential clinical applications^23^. Using predictive modeling, we therefore assessed whether the functional connectome is predictive of amyloid and tau pathology in preclinical disease. Although the relationship between functional connectivity and pathology is complex and likely bidirectional ^24^, predictive models identify statistical relationships agnostically, without requiring assumptions of directionality or causality. These identified patterns can inform hypotheses for future mechanistic studies and potentially help better detect early AD connectivity changes which may not be reflected by other modalities. We focus on the preclinical phase of AD, as it offers an opportunity to visualize tau and amyloid binding prior to widespread pathology and neuronal death; preclinical disease may also be a prime window for intervention.

We asked, does whole brain functional connectivity have predictive information for focal or global tau and amyloid PET tracer binding? To address these fundamental questions, we used connectome-based predictive modeling (CPM), a machine learning technique that identifies edges which correlate with outcomes of interest^25^. One advantage of CPM is it identifies conserved functional features across individuals, which can complement individual-specific studies of a disease process. Another major advantage is it incorporates information from the entire functional connectome, rather than limiting the scope of analysis to a specific network or region. CPM has typically been implemented to use the functional connectome as a predictor of cognitive^26–29^ and behavioral/symptomatic measures^30–33^. Here, we use CPM instead to assess to what extent the functional connectome can predict focal or global amyloid or tau binding, yielding insights into the relationship between connectivity and early-stage AD pathology.

Because tau has been shown in a growing body of literature to propagate between functionally connected regions, while amyloid’s impacted on connectivity is more global, we hypothesize that functional connectivity will better predict regional tau deposition.

## Methods

### Datasets

The Anti-Amyloid in Asymptomatic Alzheimer’s disease (A4) study is a clinical trial study of cognitively unimpaired adults (aged 65-85)^34^. Downloaded images and metrics represent baseline pre-treatment data. Specific study inclusion and exclusion criteria is described elsewhere^35^. Of those with baseline rsfMRI, 1490 participants met the requisite motion threshold (<0.3 mm mean frame-to-frame displacement) and fMRI QC (total excluded = 158). Of those with available baseline rsfMRI, 393 subjects had tau PET images (n=51 healthy controls, n = 342 amyloid-elevated individuals). Site information is unavailable for A4. We use the A4 dataset as a discovery dataset to generate our predictive models. Demographics are found in Table 1.

The Alzheimer’s Disease Neuroimaging Initiative - 3 (ADNI-3) is an observational cohort study enrolling participants from across the full spectrum of Alzheimer’s^35^. Of those with fMRI, we removed 161 participants with multi-banded fMRI due to poor data quality. 496 participants met the requisite motion threshold (<0.3 mm mean frame-to-frame displacement) and fMRI QC. Of those who were amyloid-elevated, 8 had missing tau PET metrics, leaving 211 individuals (from 45 study sites). We use the ADNI dataset an external test dataset for models generated in A4. Demographics are found in Table 1.

### Image Parameters

#### A4

Participants underwent scanning using a 3T MRI and included one resting state fMRI scan acquired through an echo-planar imaging (EPI), 2D gradient-recalled (GR), or GR EPI sequence (t = 6.5 mins, TR = 2925-3520 ms, TE = 30 ms, flip angle = 80-90°), and a T1-weighted anatomical (MPRAGE) scan (TR = 2300-7636 ms, TI = 400-900 ms, flip angle 9-11°).

Participants were imaged using GE medical systems, Philips medical systems, and Seimens scanners. Details can be found at on the A4 LONI Image and Data Archive site.

#### ADNI

Imaging consisted of a rsfMRI EPI-BOLD sequence (t = 10 min, TR/TE = 3000/30 ms, flip angle = 90°). MP-RAGE anatomical sequences were also obtained (TR = 2300ms, TI = 900 ms, flip angle = 9°). Participants were imaged using GE medical systems, Philips medical systems, and Seimens scanners. Details can be found on the ADNI LONI Image and Data Archive site.

### MRI Preprocessing

Detailed description of data preprocessing is outlined in previous work^36^. Briefly, MPRAGE scans were skull stripped using optiBET^37^ and nonlinearly aligned to the MNI-152 template using BioImage Suite (BIS^38^). Functional images were slice-time and motion corrected using SPM12, and scans with a maximum mean frame-to-frame displacement (FFD) greater than 0.3 mm were excluded to minimize motion-related artifacts^27,36^. Linear, quadratic, and cubic drifts; a 24-parameter motion model^39^; and signals from cerebrospinal fluid, white matter, and global regions were statistically removed from the data as detailed previously^40^. Connectivity matrices were generated using BIS using the Shen-268 atlas. MRIs across datasets were processed similarly. 15 ROIs were removed from connectivity matrices due to signal dropout in >50 participants (Supplementary Table 1).

### PET Preprocessing

For A4, tau PET ([18F]flortaucipir/FTP) SUVR measurements were obtained from A4, using the 90-110 minute window post-injection (4×5-minute frames). Preprocessing and analysis were performed using PETSurfer, an implementation offered in FreeSurfer (v. 6.0+). Each of the five-minute tau PET frames were motion-corrected and then summed. These composite PET images were registered to corresponding MRI images, segmented according to the Desikan-Killiany Atlas (DSK), and corrected for partial-volume effects using FreeSurfer. For each brain region defined by the DSK atlas, average tracer uptake values were calculated and standardized against the whole cerebellar cortex as the reference regions, generating standardized uptake value ratios (SUVR). For amyloid PET, global amyloid centiloid and regional amyloid SUVRs (florbetapir-PET/FBP) were obtained from the A4 online data release (Version: 2021-04-01). Amyloid metrics were derived using 50-70 minute (4×5-minute frames) post-injection images. Images were summed and realigned into a single 3D image. SUVRs were computed using the whole cerebellum as the reference region.

For ADNI, partial-volume corrected tau PET ([18F]flortaucipir/FTP) metrics were derived from the ADNI study’s records using 75-105 minute (6×5 minute frames) post-injection images.

Images were coregistered and parcellated as described above. SUVRs were computed using the inferior cerebellar cortex as the reference region.

#### Selection of ROIs

For tau analysis, we initially evaluated all 34 cortical ROIs and the amygdala in the Desikan-Killiany (DSK) atlas in the A4 cohort. To identify regions with meaningful tau accumulation, we modeled all ROIs using a two-component Gaussian Mixture Model (GMM), allowing variance parameters to vary between components and assuming the abnormal gaussian is the component with the higher mean. Regions where a two-component Gaussian distribution fit better than a one-component model (as evaluated by Bayesian Information Criterion, BIC) were assumed to have meaningful tau buildup. Of the 35 total ROIs evaluated, 26 showed superior fit with the two-component model. Upon visual inspection, we excluded 3 additional regions that deviated from the expected distributions of abnormal/elevated tau in preclinical disease. We calculated the mixing proportion of the abnormal component to estimate the number of individuals who would be considered tau-elevated in each region. ROIs with fewer than 20 estimated abnormal cases were removed to ensure sufficient signal for modeling tau patterns.

Results of this can be found in Supplementary Table 2. In addition to the 14 remaining ROIs derived in this manner, we included a temporal meta-ROI (comprised of entorhinal, parahippocampal, amygdala, fusiform, middle temporal, and inferior temporal regions) previously described in the literature^41^. For amyloid, we model an amyloid cortical composite SUVR. We also included posterior cingulate and precuneus amyloid SUVR to enable direct one-to-one comparisons with tau SUVR metrics in these same regions.

### Connectome-Based Predictive Modeling (CPM)

To identify the edges predictive of focal or global tau or amyloid in each model derived as above, we employed connectome-based predictive modeling (CPM), the specifics of which can be found in Shen *et al*^25^. Briefly, connectivity matrices (predictors) and PET/cognitive metrics (outcomes) were compiled for each participant. For each cell in the connectivity matrix (representing correlation of activity between each pair of nodes), the edge strength of all participants is aggregated and then correlated with the measures. This process was repeated for every connection in the brain, resulting in a single correlation matrix for the study, where each cell represents the correlation of the connectivity to the PET/cognitive metrics in the dataset. We use a partial correlation approach in which scanner and a mean frame-to-frame displacement are covariates when generating the study-wide correlation matrix. The connections that demonstrated a significant correlation to the outcome were selected for. For each participant, these significant connections from their connectivity matrix were summed to generate a patient-specific brain network summary score. All participants’ summary network scores were related to the outcome measure through a linear regression to model the “brain-outcome measure” relationship. The linear regression model was validated using a 5-fold cross-validation. This process was repeated 1000 times, where each time, individuals are randomly re-shuffled into new groups, generating an unbiased estimation of the study cohort.

Models were rerun with different edge-selection thresholds (p = 0.01, p = 0.05), with minimal difference in predictive power (Supplementary Table 3).

### Statistical Analyses of Models

Model performances were assessed by using a Spearman’s correlation between predicted and true PET metrics. In addition, we tested our models against permuted models, whereby the participant labels were randomly shuffled before running the model. After iterating this random permutation 1000 times, we calculated how often the accuracy of the randomly permuted predictions surpassed the median accuracy of the regular predictions. This was used to calculate a non-parametric p-value, described in Scheinost *et al*. (2021)^42^ .

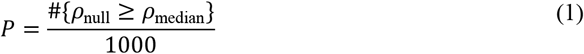

Significant models were then tested against a spatially permuted model, where each individual was assigned one of their own tau values randomly selected from the 14 individual ROIs as model input. Significance was determined as defined by equation 1. We adjusted these non-parametric p-values for multiple comparisons using the Benjamini-Hochberg procedure.

### Post-hoc analysis

Following previous studies, edges were visualized if they were found to be significant in a minimum of 4 out of 5 folds in 800 of the 1000 CPM replicates, which helps reduce noise while preserving important connections^26,31,36^. To visualize the top predictive nodes and their associated predictive edges, we generated a mask where significant correlations were represented by 1’s and non-significant edges by 0’s. From this binary mask, we calculated each node’s degree by summing the number of significant edges. We then selected the top 5% of nodes based on this degree and visualize their significant connections using circle plots and brain surfaces.

### Model Generalization

Thresholds for determination of tau negativity vs. tau-elevation were established as the SUVR value at which the posterior probability of belonging to the normal (lower mean) component exceeded 99%, as previously done^43^. We used edges that were significant in a minimum of 4 out of 5 folds in 800 of the 1000 CPM replicates within the A4 dataset. These edges, along with the corresponding beta-values from linear regression models derived from A4, were directly applied to the ADNI connectivity matrices. This process was separately applied to tau negative and tau-elevated individuals. Generalization was assessed by Spearman’s correlation of predicted and true tau PET metrics of ADNI participants.

## Results

### Functional connectivity minimally predicts local and global amyloid PET binding

We set out to determine whether functional connectivity could predict amyloid PET. We developed a model to predict global amyloid, given its use in clinical and research settings in determining amyloid positivity. We found CPM minimally predicted global amyloid (Supplementary Feig. 1; *r*_*s*_ = 0.04, p = 0.41, FDR-adjusted for 3 tests). To assess whether CPM might better predict regional amyloid SUVR over a more global measure of amyloid, we developed models to predict amyloid binding in the posterior cingulate and precuneus, regions that demonstrate the earliest buildup of amyloid in preclinical disease^44^. We did not demonstrate meaningful predictive power for amyloid binding in these regions (Supplementary Feig 1; posterior cingulate *r*_*s*_ = 0.04, p = 0.41; precuneus *r*_*s*_ = - 0.03, p = 0.60; FDR-adjusted for 3 tests).

### Functional connectivity predicts regional tau binding in preclinical AD

We next applied CPM to predict regional tau binding. CPM predicted tau SUVR in the posterior cingulate cortex, precuneus, lateral occipital, middle temporal, inferior temporal, bank STS, and superior frontal ROIs (Fig 1b, median CPM: posterior cingulate *r*_*s*_= 0.30, p<0.001; precuneus *r*_*s*_ = 0.192, p<0.001; lateral occipital *r*_*s*_ = 0.17, p=0.03; middle temporal *r*_*s*_ = 0.16, p=0.03; inferior temporal *r*_*s*_ = 0.16, p = 0.03, bank STS *r*_*s*_ = 0.15, p=0.03; superior frontal *r*_*s*_ = 0.13, p=0.03; FDR-adjusted for 15 tests), with all other ROIs not significant against permuted models. To determine whether the models that survived permutation testing contained ROI-specific information or merely reflected global tau covariance, we generated a spatially permuted model in which each individual was assigned one of their own tau values randomly selected from the 14 individual ROIs as model input. All models except the superior frontal significantly outperformed this spatially permuted model (Fig 1b, posterior cingulate p<0.001; precuneus p<0.03; lateral occipital p = 0.04; middle temporal p = 0.04; inferior temporal p = 0.04, bank STS p = 0.04, superior frontal p = 0.06, FDR-adjusted for 7 tests). When comparing the correspondence of the predictive edges between these models, we found that these tau models were spatially distinctive (Fig. 1c). There was no correlation between the prediction accuracy for a region and the estimated number of individuals in the tau-positive distribution for that region (r =-0.06). CPM tau models predicting tau z-scores (calculated relative to the 51 healthy control participants in A4) were developed to assess potential contributions of non-specific/off-target signal to prediction accuracies. These z-score models showed similar predictive accuracies as raw SUVR (Supplementary Table 4). FC versus tau and calibration plots can be found in Supplementary Feig. 2. We next assessed to what degree our tau model predictions depend on: 1) connections within the temporal lobe, including regions which have elevated tau in the earliest stages of AD (Braak I-IV) and 2) connections from the connectivity network most implicated in amnestic AD, the DMN. We first lesioned all temporal nodes, then all DMN nodes (Supplementary Feig. 3), finding that models maintained their predictive accuracies despite either lesion (Supplemental Fig. 4).

**Figure 1.**
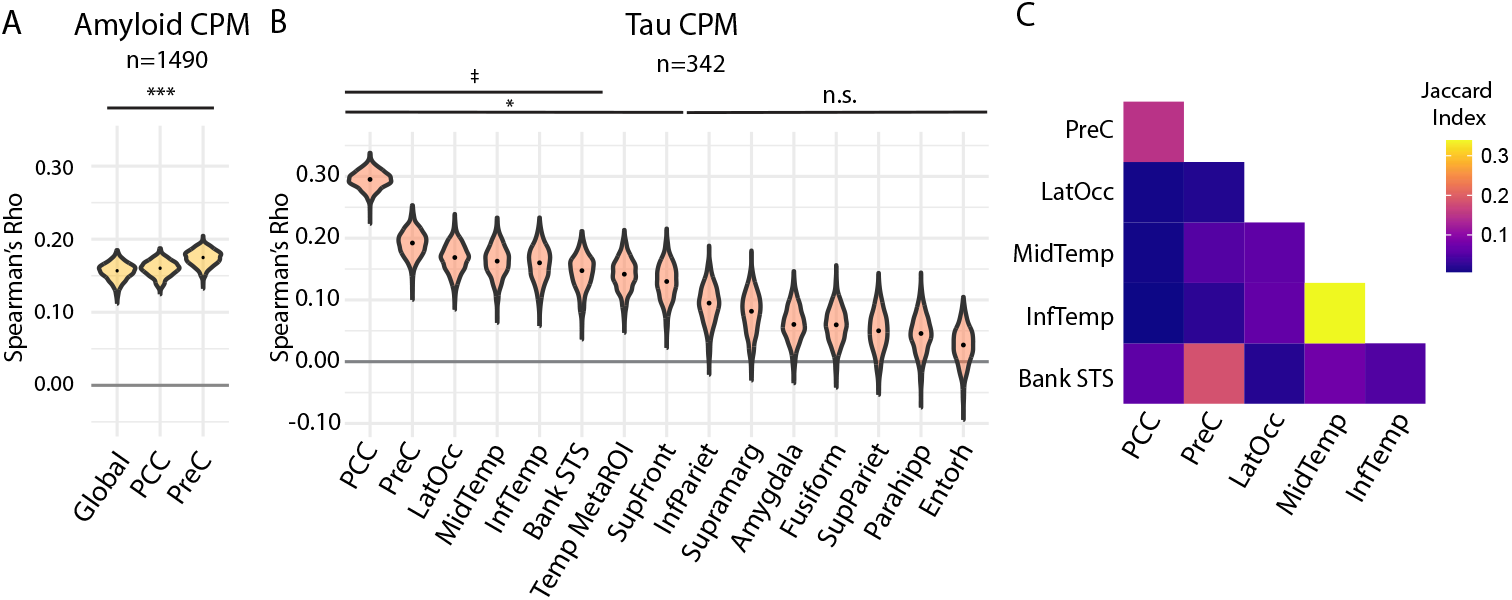
Whole brain functional connectivity predicts regional tau PET burden in preclinical Alzheimer’s disease. **(A)** Amyloid CPM results; violin plots represent distributions over 1000 model iterations of global amyloid SUVR and regional amyloid in the posterior cingulate and precuneus regions (SUVR). Model performance is measured by Spearman’s *r*_*s*_ of predicted versus observed amyloid PET. *n*=1490 amyloid positive and amyloid negative participants. *** *p*<0.001, FDR-adjusted for 3 tests (1-tailed, permutation test). **(B)** Regional tau CPM results; violin plots represent distribution over 1000 model iterations, with the median model performance depicted by the black dot. Model performance is measured by Spearman’s *r*_*s*_ of predicted versus observed regional tau PET. *n*=342 amyloid positive participants. * *p*<0.05 against permutation testing; FDR-adjusted for 14 tests (1-tailed, permutation test). ‡ p<0.05 against spatially permuted model, FDR-adjusted for 7 tests (1-tailed permutation test). **(C)** Similarity of predictive edges between regional tau models, using the Jaccard index. While this primary analysis focused on the cohort with tau PET imaging, we also generated amyloid models in the entire study population (n = 1490) to assess these findings across a full range of amyloid levels, including those below the threshold for amyloid positivity. We found these models did modestly predict amyloid levels in all 3 models (Fig. 1a; global amyloid *r*_*s*_ = 0.16, p<0.001, posterior cingulate *r*_*s*_ = 0.16, p<0.001, precuneus amyloid *r*_*s*_ = 0.17, p<0.001).

This suggests that predictors of regional tau binding are not dependent on temporal or DMN nodes.

In summary, regional tau models for posterior cingulate cortex, precuneus, lateral occipital, middle temporal, inferior temporal, and bank STS ROIs outperformed the global tau SUVR model. Additionally, when comparing equivalent regions, tau ROI models outperformed their corresponding amyloid SUVR models, despite the higher statistical power available for the amyloid models.

### Tau models generalize to an external clinical dataset (ADNI)

To assess the generalizability of our models, we sought to externally validate the 6 most predictive regional tau models to an external AD dataset, ADNI-3. ADNI-3 is a study that includes healthy controls, and those with subjective memory concerns, mild cognitive impairment, and AD dementia. Generalization was restricted to amyloid-positive individuals in ADNI (Supplementary Feig. 5). To concomitantly test model specificity, we apply models separately to tau-negative and tau-elevated individuals, defined by a 2-component GMM approach. Each of the A4 models generalized to tau-positive individuals in ADNI-3 (Fig. 2; posterior cingulate *r*_*s*_ = 0.14, Precuneus *r*_*s*_ = 0.27, lateral occipital *r*_*s*_ = 0.18, middle temporal *r*_*s*_ = 0.16, inferior temporal *r*_*s*_ = 0.14, Bank STS *r*_*s*_ = 0.25), but not to tau-negative individuals (Fig. 2; posterior cingulate *r*_*s*_ = 0.05, Precuneus *r*_*s*_ = 0., lateral occipital *r*_*s*_ = 0.00, middle temporal *r*_*s*_ = −0.18, inferior temporal *r*_*s*_ = −0.25, Bank STS *r*_*s*_ = −0.10). Given that the posterior cingulate model demonstrated the highest predictive power, we examined its underlying predictive edges and nodes in more detail. Positively correlated edges were enriched in nodes of the lateral and medial temporal lobe and cerebellum, with a particular enrichment in cross-hemispheric medial temporal connections (Fig 3a, b). Negatively correlated edges on the other hand were enriched in higher cortical association areas of the prefrontal and parietal regions (Fig. 3a, b).

**Figure 2.**
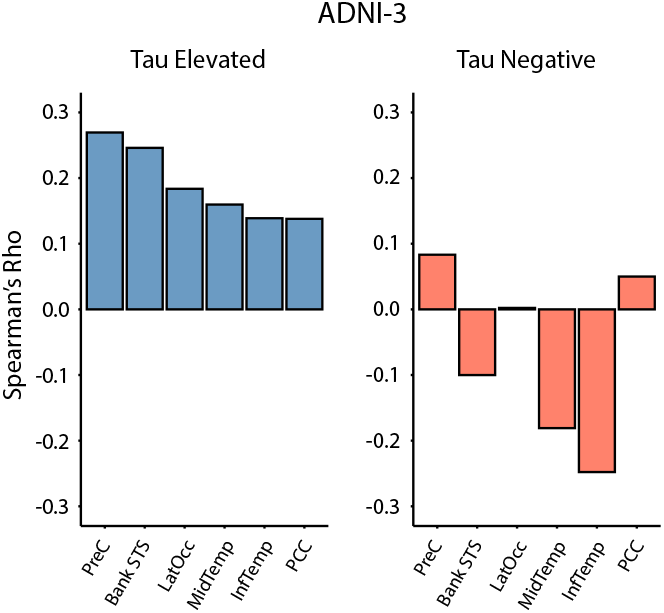
Regional tau binding models generalize to an external dataset. CPM validation of A4 regional tau models in ADNI, differentiated by tau-elevated and tau-negative groups. Model performance is measured by Spearman’s *r*_*s*_ of predicted versus observed regional tau PET in ADNI *(n* = 211).

**Figure 3.**
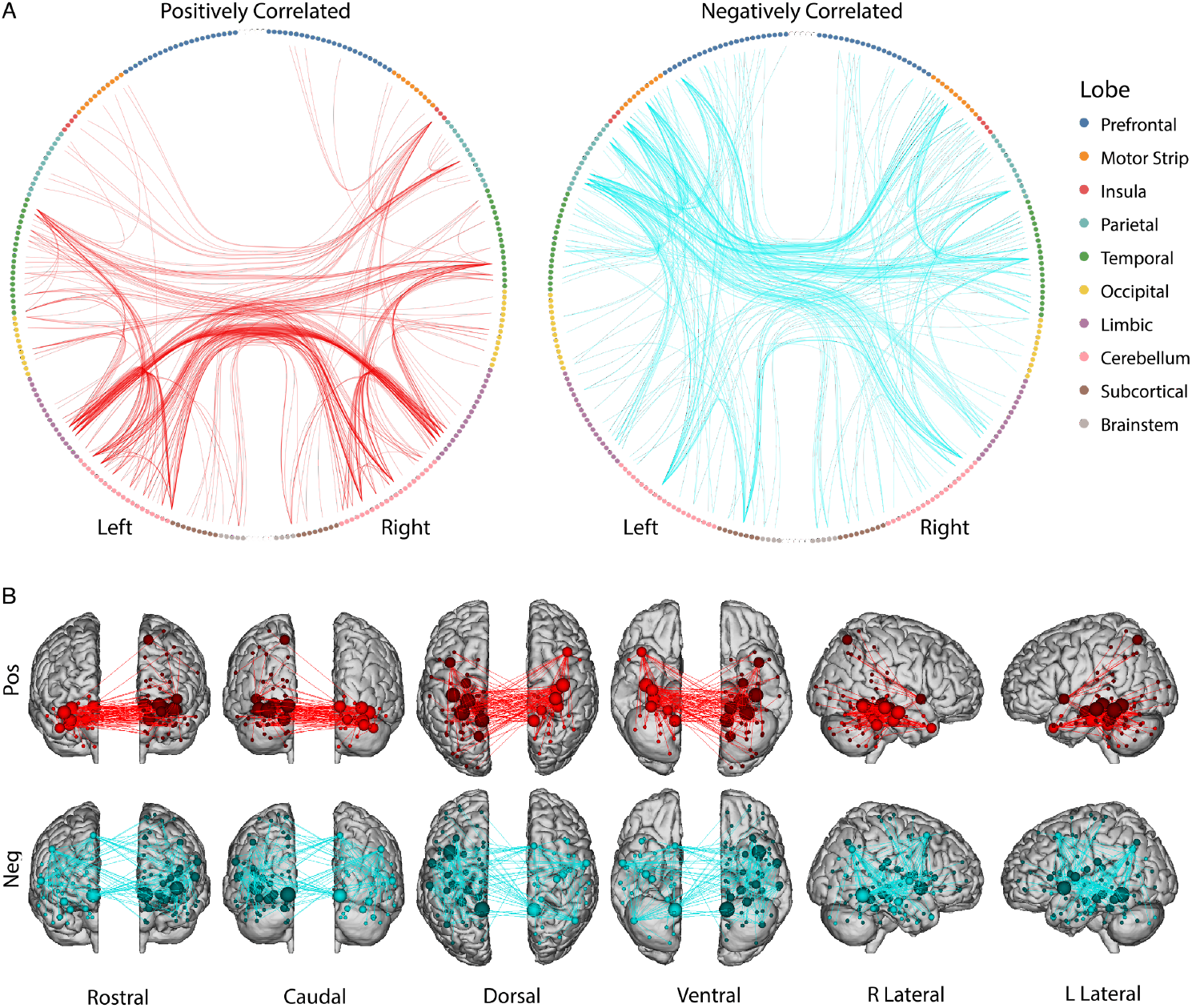
PCC tau model edges positively and negatively correlated with FC. **(A)** Circle plots of the predictive edges for the PCC model (n=342). Each circle represents a node from the Shen-268 atlas, grouped according to their respective region. Edges are depicted by lines between each node pair. Plot with red lines show edges that were positively correlated with PCC tau, while the plot with blue lines indicates edges that were negatively correlated with PCC tau. Edges displayed represent those from the top 5% most predictive nodes. **(B)** Predictive edges from circle plots plotted in (a) onto brain surfaces.

## Discussion

We use a whole-brain, data-driven approach to determine how the functional connectome predicts amyloid and tau PET binding in a preclinical AD cohort (the A4 Study). We demonstrate that functional connectivity is a predictor of regional tau binding, but a poorer predictor of amyloid binding. Tau models were most accurate for Braak IV/V regions (posterior cingulate, precuneus, lateral occipital cortex, middle temporal, inferior temporal, and Bank STS). These models generalized to an external dataset (ADNI-3) including both symptomatic and preclinical individuals with AD. Notably, models generalized specifically to tau-positive and not to tau-negative members of the ADNI cohort, emphasizing these models are not meaningful in individuals without abnormal tau. The underlying predictive edges of the PCC model (the most accurate in A4) suggest that there is a spatial distinction between the most predictive positively correlated and negatively correlated nodes and their respective edges.

Our use of CPM represents a novel direction from its typical use. While CPM has traditionally been used to relate connectome patterns to behavioral and psychiatric measures, we uniquely apply it to predict well-defined, biologically validated metrics of early-stage AD pathology.

Multimodal predictive modeling, especially where one modality is well-described and well-validated, allows for investigations of functional connectivity to be anchored in biologically informed ways. That is, because tau and amyloid PET are reliable and valid measures of underlying AD pathology, we can use PET as an indirect way to more accurately, and agnostically, assess connectivity differences that track with AD progression. Furthermore, by incorporating additional dimensions to our analyses, multimodal approaches may enhance sensitivity to uncover disease characteristics not discernible from unimodal analyses, which may help better identify clinically meaningful subgroups. With interpretable models such as CPM, wherein one can identify the edges and nodes predictive of the outcome, we can also generate new hypotheses to test in future mechanistic studies.

A key finding of this work was that the accuracy of our models in predicting regional tau outperformed models predicting regional and global amyloid. Recent work has demonstrated an association between whole-brain tau PET covariance and functional connectivity^8,45^. We add to this existing literature by demonstrating functional connectivity can predict regional tau PET signal, but minimally predicts amyloid PET. This finding converges with demonstrations that tau progresses along pathways of functional connectivity^7–9^, and is more closely associated than amyloid with disease outcomes such as cortical atrophy and symptomatology^46–48^. We demonstrate that regional tau models convey meaningful region-specific information that is not captured by global tau covariance, outperforming spatially permuted models. Regional tau models also outperform the temporal meta-ROI model, suggesting that there is meaningful variation in regional tau’s relationship to functional connectivity that is lost in a meta-ROI approach. Future work should examine how these models improve using functional parcellations of tau-PET, which would allow 1:1 comparisons of tau PET to FC ROIs.

Interestingly our models were more accurate in predicting tau for regions associated with Braak stages (IV-VI) than those corresponding to earlier Braak stages (I/III). One interpretation is that functional connectivity in later-stage regions may be particularly sensitive to tau deposition, such that even modest tau buildup may lead to disproportionate FC disruptions. Alternatively, tau deposition might be driven by FC shifts in these regions, which are known to participate in major networks (e.g., PCC and precuneus of the DMN). Recent work applying CPM to familial AD (fAD) corroborates our finding that precuneus tau can be predicted from functional connectivity^49^. However, unlike our preclinical AD models, this study also found high predictability of entorhinal cortex tau – a difference that may reflect the distinct disease trajectories between sporadic and genetic AD patients.

When examining the underlying predictive edges for the PCC model, we saw a mixture of positively and negatively correlated edges associated with regional tau SUVR; positively correlated edges were enriched in nodes of temporal lobe, limbic, and cerebellar regions, whereas negatively correlated edges were enriched in nodes of heteromodal association cortices such as the prefrontal and parietal regions. It is possible that edges positively correlated with regional PCC tau represent compensatory connectivity, whereas edges negatively correlated with regional PCC tau represent a loss of normal connectivity. However, more work will be needed to more precisely distill these associations. Future work should also establish whether these identified edges may inform our ability able to identify at-risk, clinically meaningful subgroups in preclinical disease.

While this work was able to generate generalizable models in preclinical AD from a single six-minute resting state fMRI scan, future datasets which incorporate longer scan times and continuous performance task-based fMRI, or which take advantage of second-generation PET tracers and next-generation PET technology for finer resolution, could improve connectivity estimates and prediction performance^27,50^. There are other caveats to the current work. The A4 study restricted enrollment to individuals who performed within 1.5 s.d. of the normative mean on a test of short-term memory^34^. This may impact model generalizability in participants whose cognitive measures deviate from the average. Encouragingly, however, our results do generalize to an external dataset with individuals with a broader range of clinical disease and cognitive test performance, suggesting they may be relevant regardless of cognitive performance. This highlights the utility of neuroimaging to detect and track preclinical AD pathophysiology, even in the absence of overt cognitive changes. Functional connectivity may be an important complement to cognitive screening and plasma/CSF biomarkers in predicting trajectories in preclinical AD.

While CPM can help generate new hypotheses to better understand the disease, the ability to define causal relationships with predictive models is limited. Importantly, this predictive modeling approach does not imply unidirectional relationship between functional connectivity and tau pathology. Our findings should therefore be further explored through quasi-experimental statistical techniques and animal experiments^51^. For example, one such quasi-experimental technique, regression discontinuity, could be used to explore connectivity changes above and below certain critical tau thresholds in individuals given amyloid/tau clearing agents, helping to determine the causal effects of AD pathology on connectivity. A difference-in-differences approach could compare the change in functional connectivity over time in immediately adjacent regions with high and low tau accumulation, estimating the causal impact of tau on connectivity changes. Animal experiments can also help causally assess some of the hypotheses we pose here. For instance, to determine whether the positively correlated edges we identify here are compensatory (Fig. 3a, b), researchers could study dose-dependent concentrations of tau in murine temporal ROIs while simultaneously using optogenetic inhibition of these ROIs during a memory task; if they are compensatory, the group with inhibition would experience earlier task-associated deficits compared to the control group.

It will also be vital to more precisely assess the temporal aspects of disease progression. Past work has shown models defined across an entire sample might not be relevant for all individuals in the dataset^52^. This is especially important in AD, which exhibits significant spatiotemporal heterogeneity. While amyloid-positive individuals are at high risk for future AD development, asymptomatic amyloid positivity is a large window, with the first positive test about 10-20 years prior to symptom onset^11^. Better defining clinically meaningful subgroups within this preclinical stage may aid in future risk assessment and treatment selection. Techniques such as the subtype and stage inference algorithm (SuStain), which pseudotemporally assigns individuals into different stages of the disease while simultaneously clustering individuals based on similarities in disease progression^53,54^, may help in this effort. Future longitudinal studies can also help distinguish temporal heterogeneity of individual patients across all forms of AD. The emergence of large preclinical datasets such as A4 represent a significant milestone in the field, allowing us to use robust statistical techniques to model preclinical illness for the first time. The potential to identify individuals with preclinical disease using highly sensitive and specific blood-based biomarkers will aid immensely in the early identification and, ultimately, treatment of AD^55^.

Our results underscore the ability of the functional connectome to predict regional tau deposition even in the preclinical stages of AD. It is striking that we can best predict tau PET binding in highly interconnected nodes that are associated with the DMN and with Braak stages IV and V pathology, such as the posterior cingulate, precuneus, and lateral occipital regions. We hypothesize that this suggests major brain hubs (some of the strongest of which are part of the DMN) may play a critical role in the dissemination of tau in preclinical disease. Combining advanced neuroimaging techniques with computational approaches has the potential to identify AD progression, classify individuals into AD subgroups, and potentially reveal new therapeutic targets for personalized intervention in the earliest stages of illness.

## Data Availability

Data from the A4 and ADNI-3 studies can be requested online.

## Acknowledgements

The A4 and LEARN Studies were led by Dr. Reisa Sperling and Dr. Paul Aisen. The complete A4 Study Team list is available on: https://www.actcinfo.org/a4-study-team-lists/. We would like to acknowledge the dedication of the study participants and their study partners who made the A4 and LEARN studies possible.

## Supplementary material

Supplementary material is available online.

## Code availability

Code for CPM scripts are available at https://github.com/orgs/frederickslab/repositories

