## Supplementary Information for "Whole-brain functional connectivity predicts regional tau PET in preclinical Alzheimer’s disease"

Supplemental Material

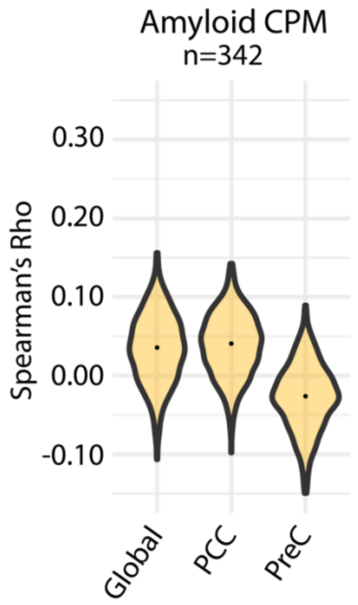


Supplemental Figure 1. Amyloid CPM models for tau PET cohort (n = 342)

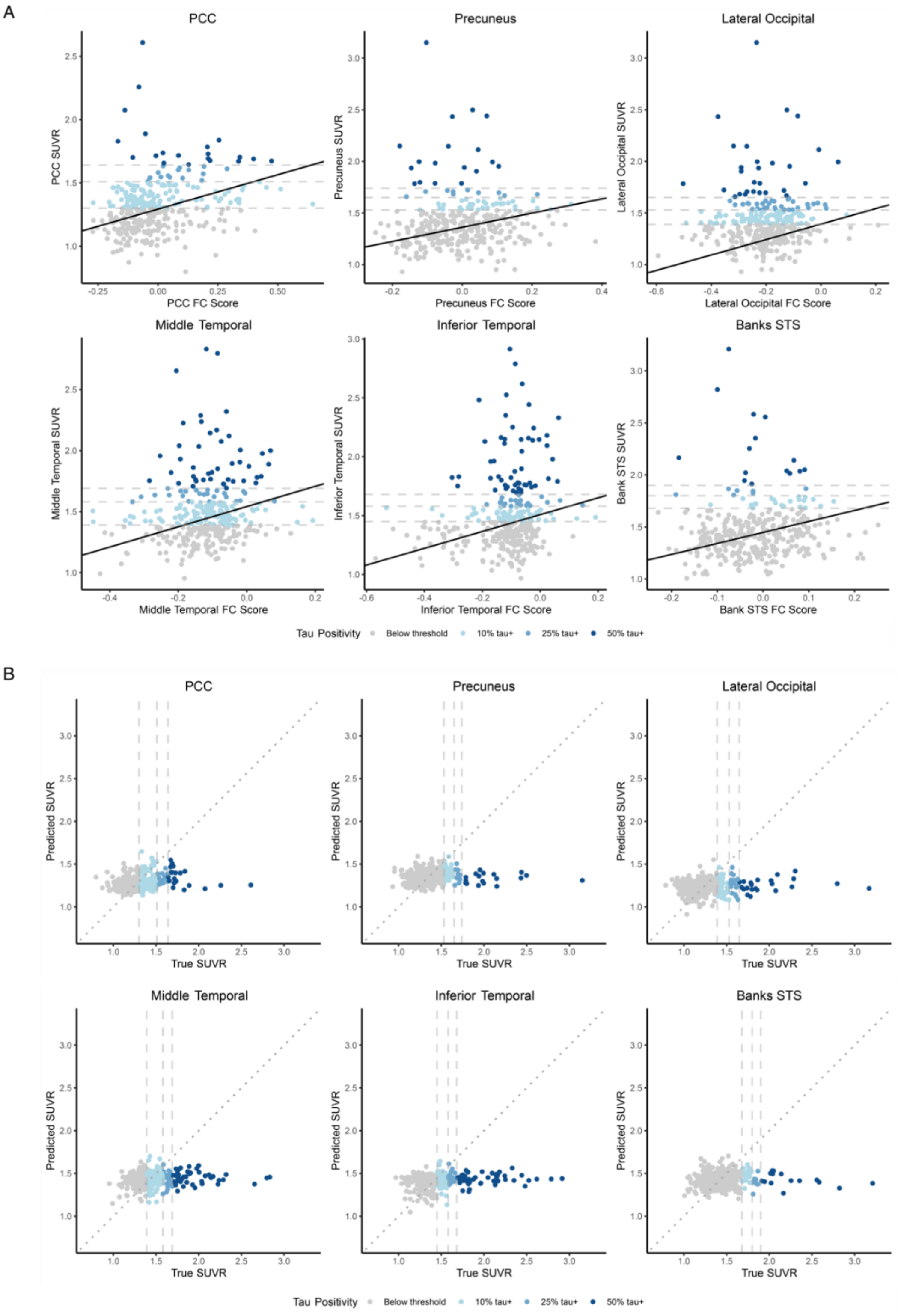


Supplemental Figure 2. A) Plots of FC score vs ROI SUVR for significant models. Lines defined by regression parameters from CPM (n=342). B) Calibration plots of true SUVR vs predicted SUVR. For both panels, tau positivity determined by posterior probability of belonging to abnormal component and are colored for various thresholds.

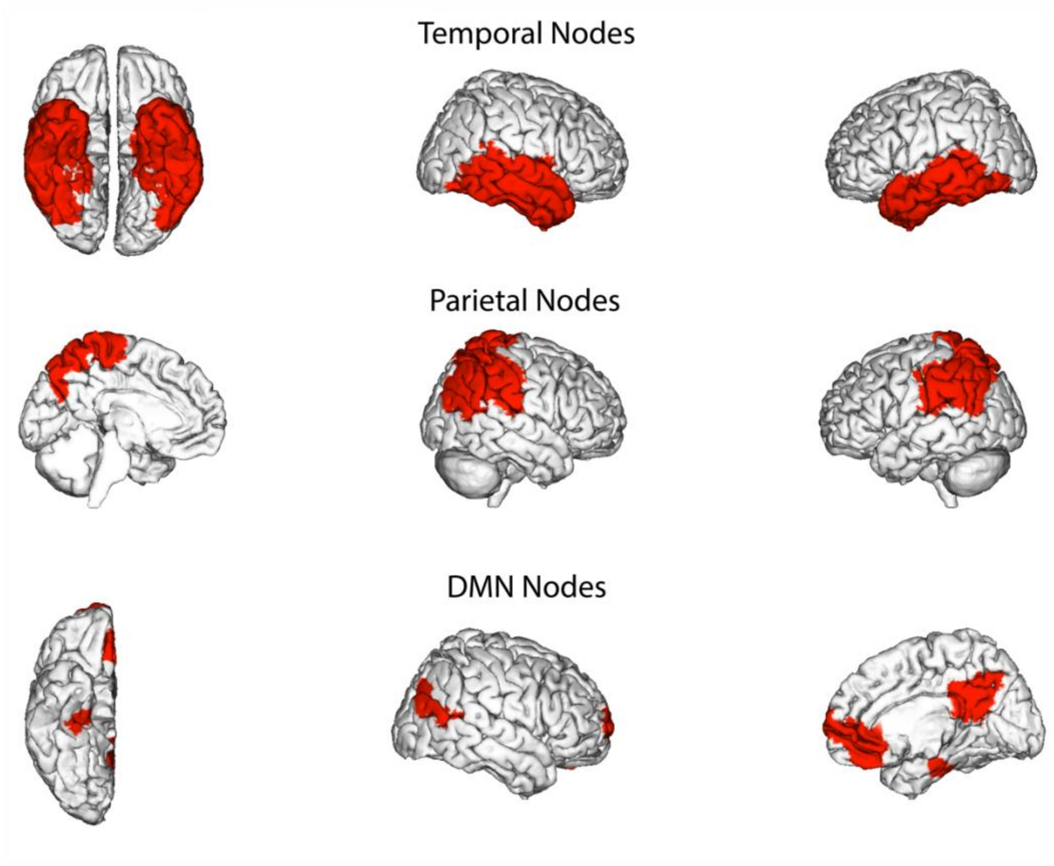


Supplemental Figure 3. Visualization of nodes defined as temporal lobe, parietal lobe, and DMN.

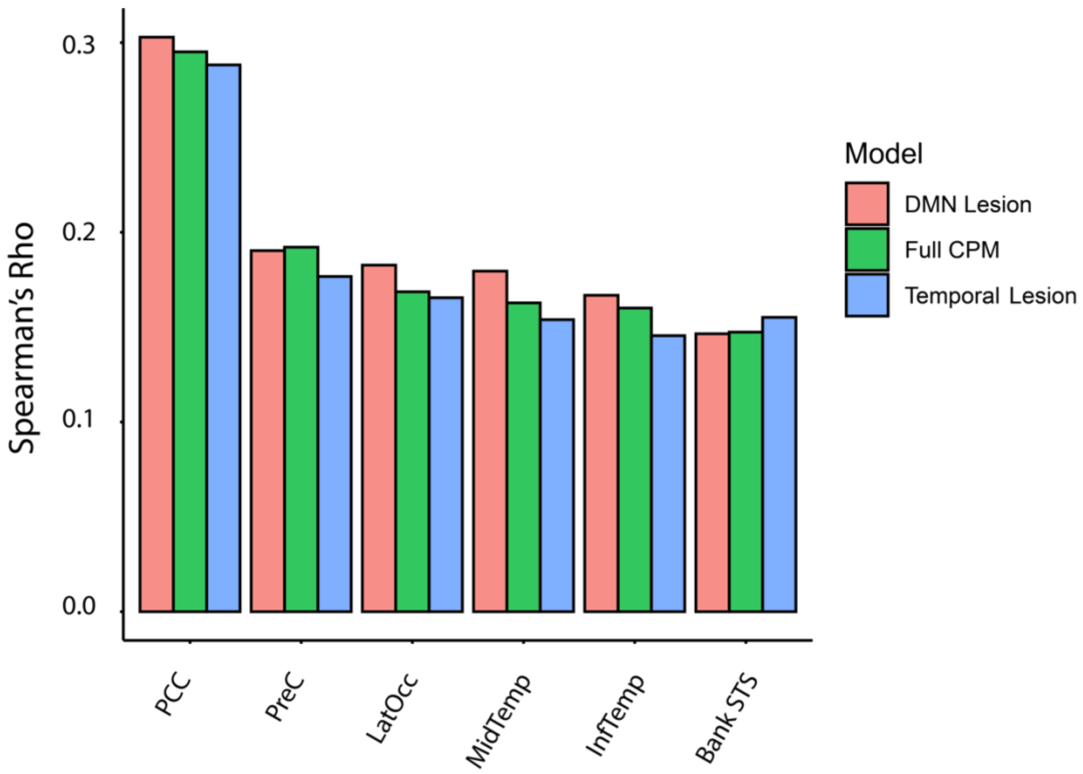


Supplemental Figure 4. Predictive accuracies of tau models when lesioning of the temporal lobe and DMN-associated nodes.

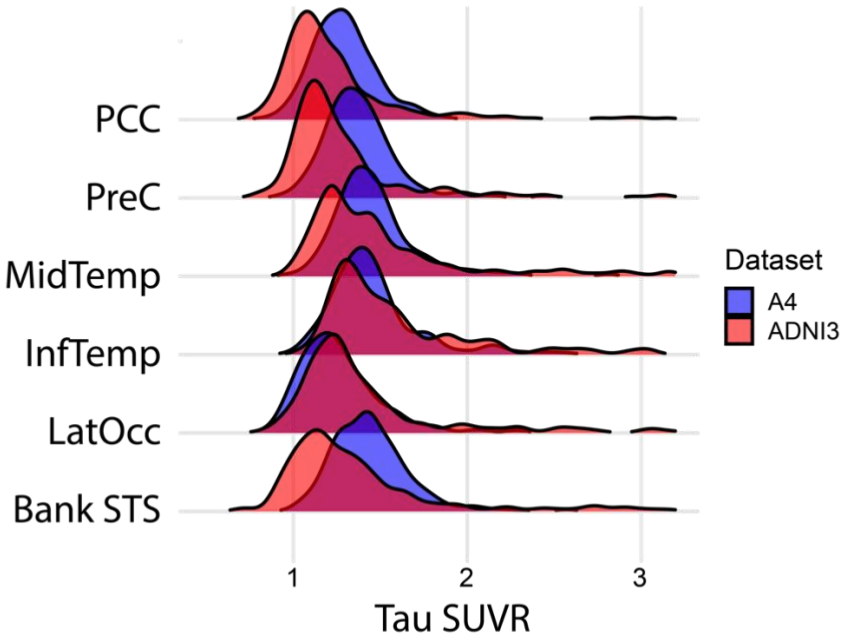


Supplemental Figure 5. Visualization of tau SUVR distributions in A4 and ADNI3 in amyloid-positive participants.

| NODE (SHEN ATLAS) | # PT MISSING | TAL X | TAL Y | TAL Z | MNIX | MNIY | MNIZ | BA REGION |
|---|---|---|---|---|---|---|---|---|
| **2** | 200 | 9 | 14 | -14 | 10 | 18 | -19 | Right-OrbFrontal (11) |
| **4** | 247 | 15 | 29 | -20 | 16 | 34 | -23 | Right-OrbFrontal (11) |
| **51** | 130 | 26 | 6 | -31 | 27 | 12 | -40 | Right-Temporalpole (38) |
| **60** | 127 | 30 | -4 | -35 | 31 | 0 | -45 | Right-InfTempGyrus (20) |
| **107** | 58 | 45 | -49 | 0 | 46 | -49 | -6 | Right-Fusiform (37) |
| **131** | 157 | 5 | -24 | -32 | 6 | -22 | -42 | Outside defined Bas |
| **135** | 87 | -18 | 14 | -16 | -18 | 19 | -21 | Left-OrbFrontal (11) |
| **136** | 518 | -6 | 14 | -18 | -6 | 18 | -23 | Left-OrbFrontal (11) |
| **137** | 104 | -8 | 34 | -20 | -8 | 39 | -21 | Left-OrbFrontal (11) |
| **189** | 282 | -23 | 4 | -31 | -23 | 9 | -39 | Left-Temporalpole (38) |
| **196** | 150 | -49 | -20 | -22 | -52 | -18 | -29 | Left-InfTempGyrus (20) |
| **202** | 145 | -29 | -9 | -32 | -30 | -6 | -41 | Left-Parahipp (36) |
| **249** | 52 | -34 | -53 | -43 | -35 | -50 | -54 | Outside defined BAs |
| **252** | 120 | -44 | -50 | -34 | -46 | -47 | -44 | Outside defined BAs |
| **268** | 111 | -5 | -21 | -27 | -7 | -19 | -37 | Outside defined BAs |

Supplementary Table 1. List of nodes with greater than 50 participants (of the total 1490 participants) with signal dropout. Nodes listed above were removed from connectivity matrices prior to all analyses.

| ROI | BIC<br><br>1-component | BIC<br><br>2-component | Delta BIC | Abnormal Component mixing proportion, (estimated n) |
|---|---|---|---|---|
| Inferior Parietal | -319 | -448 | -129 | 0.133 (45) |
| Fusiform | -455 | -550 | -95 | 0.172 (59) |
| Transverse Temporal | 144 | 54.6 | -89.4 | NA |
| Inferior Temporal | -418 | -501 | -83 | 0.184 (63) |
| Rostral Middle Frontal | -284 | -361 | -77 | 0.027 (9) |
| Superior Parietal | -308 | -382 | -74 | 0.079 (27) |
| Isthmus Cingulate | -282 | -356 | -74 | 0.055 (18) |
| Caudal Middle Fontal | -240 | -303 | -63 | 0.039 (13) |
| Precuneus | -372 | -434 | -62 | 0.081 (27) |
| Middle Temporal | -378 | -431 | -53 | 0.173 (59) |
| Bank STS | -361 | -405 | -44 | 0.075 (25) |
| Lateral Occipital | -346 | -387 | -41 | 0.122 (41) |
| Amygdala | -222 | -260 | -38 | 0.241 (82) |
| Supramarginal | -50.1 | -86.3 | -36.2 | 0.137 (47) |
| Superior Frontal | -488 | -519 | -31 | 0.105 (36) |
| Paracentral | -385 | -405 | -20 | 0.017 (6) |
| Frontal Pole | -165 | -182 | -17 | NA |
| Cuneus | 273 | 262 | -11 | 0.022 (7) |
| Parstriangularis | -476 | -486 | -10 | NA |
| Posterior Cingulate | -441 | -448 | -7 | 0.123 (42) |
| Parahippocampal | -380 | -387 | -7 | 0.065 (22) |
| Lingual | -148 | -154 | -6 | 0.046 (16) |
| Precentral | -515 | -520 | -5 | 0.022 (8) |
| Parsorbitalis | -542 | -546 | -4 | 0.037 (13) |
| Entorhinal | -291 | -295 | -4 | 0.111 (40) |

| | | | | |
|---|---|---|---|---|
| Parsopercularis | -17.5 | -18.5 | -1 | 0.02 (7) |
| Temporal Pole | -434 | -434 | 0 | NA |
| Rostral Anterior Cingulate | -273 | -273 | 0 | NA |
| Postcentral | -294 | -293 | 1 | NA |
| Caudal Anterior Cingulate | -441 | -437 | 4 | NA |
| Superior Temporal | -258 | -251 | 7 | NA |
| Medial Orbitofrontal | -590 | -583 | 7 | NA |
| Lateral Orbitofrontal | -404 | -395 | 9 | NA |
| Pericalcarine | -454 | -444 | 10 | NA |
| Insula | -351 | -336 | 15 | NA |

Supplementary Table 2. For each region of interest (ROI) from A4 (n=342), the table reports the BIC of the one-component and two-component Gaussian mixture models, along with the delta BIC (2-component – 1 component) and mixing proportion of the abnormal/elevated tau component. We do not report mixing proportions for models where 1-component>2-component, and for regions with abnormal 2-component GMM: pars triangularis (due to approximately equal Gaussian means between components), the transverse temporal cortex (where the abnormal component AUC exceeded the normal component), and frontal pole (abnormally high mixing proportion of the abnormal component).

| ROI | Spearman's Rho (edge p = 0.05) | Spearman's Rho (edge p = 0.01) |
|---|---|---|
| Posterior Cingulate | 0.295 | 0.282 |
| Precuneus | 0.192 | 0.186 |
| Lateral Occipital | 0.169 | 0.155 |
| Middle Temporal | 0.163 | 0.153 |
| Inferior Temporal | 0.160 | 0.142 |
| Bank STS | 0.147 | 0.134 |
| Temporal -ROI | 0.142 | 0.121 |
| Superior Frontal | 0.130 | 0.142 |
| Inferior Parietal | 0.095 | 0.103 |
| Supramarginal | 0.082 | 0.087 |
| Amygdala | 0.060 | 0.040 |
| Fusiform | 0.060 | 0.059 |
| Superior Parietal | 0.050 | 0.050 |
| Parahippocampal | 0.046 | 0.045 |
| Entorhinal | 0.027 | 0.028 |

Supplementary Table 3. Comparison of model performance metrics between models with feature selection threshold of p = 0.05 and p = 0.01 for n = 342 subjects (amyloid elevated only).

| ROI | Median Spearman’s rho |
|---|---|
| Posterior Cingulate | 0.293 |
| Precuneus | 0.193 |
| Lateral Occipital | 0.170 |
| Middle Temporal | 0.167 |
| Inferior Temporal | 0.161 |
| Bank STS | 0.150 |

Supplementary Table 4. Model performance (median spearman’s) using z-scored tau PET SUVRs as model inputs.
